# Sex differences in the distribution and density of regulatory interneurons in the striatum

**DOI:** 10.1101/2024.02.29.582798

**Authors:** Meghan Van Zandt, Deirdre Flanagan, Christopher Pittenger

## Abstract

Dysfunction of the cortico-basal circuitry – including its primary input nucleus, the striatum – contributes to neuropsychiatric disorders, including autism and Tourette Syndrome (TS). These conditions show marked sex differences, occurring more often in males than in females. Regulatory interneurons, including cholinergic interneurons (CINs) and parvalbumin-expressing GABAergic fast spiking interneurons (FSIs), are implicated in human neuropsychiatric disorders such as TS, and ablation of these interneurons produces relevant behavioral pathology in male mice, but not in females. Here we investigate sex differences in the density and distribution of striatal interneurons, using stereological quantification of CINs, FSIs, and somatostatin-expressing (SOM) GABAergic interneurons in the dorsal striatum (caudate-putamen) and the ventral striatum (nucleus accumbens) in male and female mice. Males have a higher density of CINs than females, especially in the dorsal striatum; females have equal distribution between dorsal and ventral striatum. FSIs showed similar effects, with a greater dorsal-ventral density gradient in males than in females. SOM interneurons were denser in the ventral than in the dorsal striatum, with no sex differences. These sex differences in the density and distribution of FSIs and CINs may contribute to sex differences in basal ganglia function, including in the context of psychopathology.

## Introduction

The cortico-basal circuitry, particularly the striatum (caudate, putamen, and nucleus accumbens), is a locus of pathology in numerous neuropsychiatric disorders, including autism and Tourette’s Syndrome (TS) (Chen et al, 2022; Macpherson & Hikida, 2019; Swerdlow & Koob, 1987). The striatum is the primary input nucleus and a central hub of the basal ganglia circuitry; it is implicated in numerous neural processes, including motion, effort, habit formation, motivation, and social behavior (Bamford & Bamford, 2019; Graybiel et al, 1994; Graybiel & Grafton, 2015; Pennartz et al, 2009; Suzuki et al, 2021).

The striatum is composed of GABAergic medium spiny neurons (MSNs, comprising ∼95% of the neurons in the striatum in rodents)(Gerfen & Wilson, 1996) alongside several populations of regulatory interneurons. These include ChAT-expressing cholinergic interneurons (CINs) and multiple subtypes of GABAergic interneurons, typically identified by their expression of markers such as parvalbumin (PV) and somatostatin (SOM) (Kawaguchi, 1997; Kawaguchi et al, 1995; Marin et al, 2000; Muñoz-Manchado et al, 2018). CINs innervate various GABAergic interneurons, including PV-expressing fast-spiking interneurons (FSIs) (Chang & Kita, 1992; Kocaturk et al, 2022). They also reciprocally regulate dopamine release (Chantranupong et al, 2023; Howe et al, 2019; Yorgason et al, 2017) and can regulate excitatory glutamatergic input to MSNs (Ian Antón & Jun, 2011; Pakhotin & Bracci, 2007). FSIs are important regulators responsible for fine tuning MSN firing (Bennett & Bolam, 1994; Orduz et al, 2013) and integrate with other interneurons in coordinating movement bouts (Gritton et al, 2019) and regulating behavior (Lee et al, 2017). SOM interneurons play a complex role in regulating MSNs, FSIs, and CINs (Cattaneo et al, 2019; Momiyama & Zaborszky, 2006) and interact with dopaminergic afferents (Gazan et al, 2020; Hathway et al, 1999). Despite being a small percentage of striatal neurons, these interneurons critically regulate striatal function and behavioral output.

Interneuron pathology has been implicated in numerous neuropsychiatric disorders (Brady et al, 2022; Kataoka et al, 2010; Poppi et al, 2021; Rapanelli et al, 2017a). Deficits in cholinergic function are implicated in Parkinson’s, Huntington’s, and Alzheimer’s Diseases, schizophrenia, bipolar disorder, and attention deficit-hyperactivity disorder (Lim et al, 2014). Postmortem studies in human patients with TS have shown significant reductions in striatal cholinergic (CINs) and parvalbumin-expressing (PV) GABAergic interneurons (Kalanithi et al, 2005; Kataoka et al, 2010; Lennington et al, 2016).

Many of these conditions affect males and females differentially; for example, TS is diagnosed approximately twice as often in males as in females (Martino et al, 2013; Scahill et al, 2014). Whether striatal interneuron pathology is similarly sexually dimorphic is unknown, but preclinical evidence is beginning to suggest that it may be. The striatal circuitry can be impacted by the estrous cycle (Cao et al, 2018; Zachry et al, 2021); both GABAergic and cholinergic interneurons express estrogen receptors (Almey et al, 2012; Almey et al, 2016). Furthermore, depletion of CINs during development (Cadeddu et al, 2023), or conjoint depletion of both CINs and FSIs in adults (Rapanelli et al, 2017b), produces dysregulated striatal activity and behavioral abnormalities of potential relevance to TS and related conditions – repetitive behavioral pathology, anxiety, and social deficits – in male mice, but not in females. This suggests an underlying sex difference in striatal interneurons, their regulation of local microcircuits, and their role in the modulation of striatum-dependent behaviors.

Sex differences have been described in several aspects of the striatal circuitry. MSNs exhibit significantly higher density and size of dendritic spines in females than in males (Forlano & Woolley, 2010); estrogen modulates spine density (Peterson et al, 2015; Staffend et al, 2011). MSN excitability varies with estrous phase, with higher spontaneous EPSC frequencies during the estrous and proestrus phases (Aziz & Mangieri, 2023; Proaño et al, 2018). Endogenous striatal dopamine levels flux with the estrous cycle (Castner et al, 1993; Xiao & Becker, 1994); estrogen has been shown to potentiate dopamine release (Becker & Beer, 1986; Yoest et al, 2018), and females have higher striatal dopamine release and cycling (Calipari et al, 2017; Dluzen & McDermott, 2008; Walker et al, 2000; Zachry et al, 2021). However, whether these sex differences extend to regulatory interneurons of the striatum, such as CINs and FSIs, is not yet clear.

Here, using mice as a model system, we provide the first rigorous comparison between males and females of the density and distribution of three different subtypes of striatal interneurons implicated in striatal function and in neuropsychiatric disease: PV- and SOM-expressing GABAergic interneurons and CINs.

## Materials and Methods

### Experimental Design

All animal use followed protocols approved by Yale’s Institutional Animal Care and Use Committee. Adult male and female wild-type C57BL/6 mice (10-14 weeks old) were group housed in our animal facility in self-ventilating cages, maintained on a 12hr light/dark cycle, and provided with food and water *ad libitium*.

For histology, mice were anesthetized (ketamine 100mg/kg and xylazine 10mg/kg, followed by isoflurane inhalation until no response to noxious stimulus) followed by transcardial perfusion with 4% paraformaldehyde in phosphate buffered saline (PBS) (Thermo Fisher) and brain extraction. Cryostat sections were cut at 30 μm and mounted on Diamond White Glass Slides (Globe). For immunostaining for identification and counting of interneurons, slides were incubated overnight at room temperature with primary antibodies diluted in 0.1 M PBS containing 10% normal goat serum (Vector Labs) and 1% Triton-X 100 (AmericanBio). Primary antibodies were used as follows: Rabbit recombinant monoclonal anti-somatostatin (1:250, Abcam, ab111912), mouse monoclonal anti-ChAT (1:250, Thermo Fisher, MA5-31383), rabbit polyclonal anti-PV (1:250, Abcam, ab11427). Detection was performed with appropriate secondary antibodies: polyclonal goat anti-rabbit coupled to Alexa Fluor 488 (1:1000, Thermo Fisher, A-11008), polyclonal goat anti-rabbit coupled to Alexa Fluor 568 (1:1000, Thermo Fisher, A-11011), recombinant polyclonal goat anti-mouse coupled to Alexa Fluor 488 (Thermo Fisher, A-28175), and polyclonal goat anti-mouse coupled to Alexa Fluor 568 (1:1000, Thermo Fisher, A-11004). Sections were then mounted and coverslipped in ProLong Gold with DAPI (Life Technologies).

Immunostained coronal striatal sections (30 μm, bregma 1.1mm-0.50mm) were visualized on a Zeiss Scope.A1 using a Plan-APOCHROMAT 10x objective and an Axiocam 503 Mono at 100x magnification. Microscope fields were systematically tiled over the entire dorsal and ventral striatum and stitched together using the stitching plugin in FIJI (Schindelin et al, 2012) to provide a high-resolution image throughout the extent of the structure. Only undamaged sections with consistent immunostaining through the full extent of the striatum were included in analysis. Sections were analyzed blind to sex. Minor variances in background staining can occur with image stitching, but these did not interfere with quantification as stained neurons remained well-defined and distinct from background.

Whole striatal slice reconstructions were imported into StereoInvestigator 10 (MBF Biosciences) and regions of interest were defined. Sections were selected from the anterior striatum, between bregma 1.10-0.50mm, where the NAcc is well represented. For whole striatal analyses, the entire anatomical area of the striatum contained in the image, bounded by the internal capsule/corpus callosum white matter and other landmarks following the atlas of Paxinos (Franklin & Paxinos, 1997), was selected for quantification. For dorsal/ventral subregion analyses we identified caudate-putamen (CPu) and nucleus accumbens (NAcc) following Franklin and Paxinos (1997) and separately quantified interneuron density in these subregions. To further characterize dorsal-ventral differences in interneurons density, in a secondary analysis the striatum was divided in half by a horizontal line placed halfway between the dorsal and ventral extent of the structure in each slice (see Supplemental Figure 1A-C); quantification of interneurons using this arbitrary division of the striatum is presented in the Supplement. Neurons were counted using the fractionator tool in StereoInvestigator using a counting box size of 100 x 100 μm, excluding all neurons on the dashed bounding line. Only neurons with clearly defined soma distinct from background were counted. Data collected from left and right striatum were averaged. In total, 22 male and 17 female brains were analyzed, with two randomly selected sections per animal from within the range specified above.

### Statistics

All data were imported into JASP Statistical Software (Eric-Jan Wagenmakers, University of Amsterdam) for statistical analysis. Data was tested for normality using the D’Agostino & Pearson test and outliers were excluded. For direct comparisons between sexes for whole striatum and to compare ratios of dorsal to ventral interneurons, a two-tailed unpaired t-test was performed. To compare between sexes for the dorsal and ventral sub-regions of the striatum, a two-way repeated-measures ANOVA was used, followed by Fisher’s LSD post-hoc test (Table One). Graphs were generated using GraphPad Prism 10 (Graph Pad Software, LLC). Whiskers on bar and whisker plots indicate the min and max values, with all data points represented on graphs.

## Results

We first quantified CIN density in male and female mice (Figure 1A-D). Male mice had a significantly higher density of ChAT+ interneurons in the whole striatum (Figure 1E; two-tailed t-test, t[26] = 2.305, p=0.031). Males and females showed different patterns of CIN distribution in in anatomically defined subregions, the CPu and NAcc. There were significant effects of both sex (Figure 1F; Repeated Measures ANOVA: F(1, 28) = 4.769, p = 0.037) and subregion (F(1, 28) = 5.991= 5.991, p=0.021), with no interaction (sex x subregion F(1, 28) = 0.6137, p=0.44). The density of ChAT interneurons in the female striatum was not different in the CPu and the NAcc (Fisher’s LSD, p=0.278). In contrast, males showed significantly higher density of ChAT+ neurons in the CPu compared to the NAcc (Fisher’s LSD: p=0.026). Males had higher overall CIN density in the CPu than females (Fisher’s LSD: p=0.021), but there were no differences in the NAcc (Fisher’s LSD: p=0.279).

**Figure 1.**
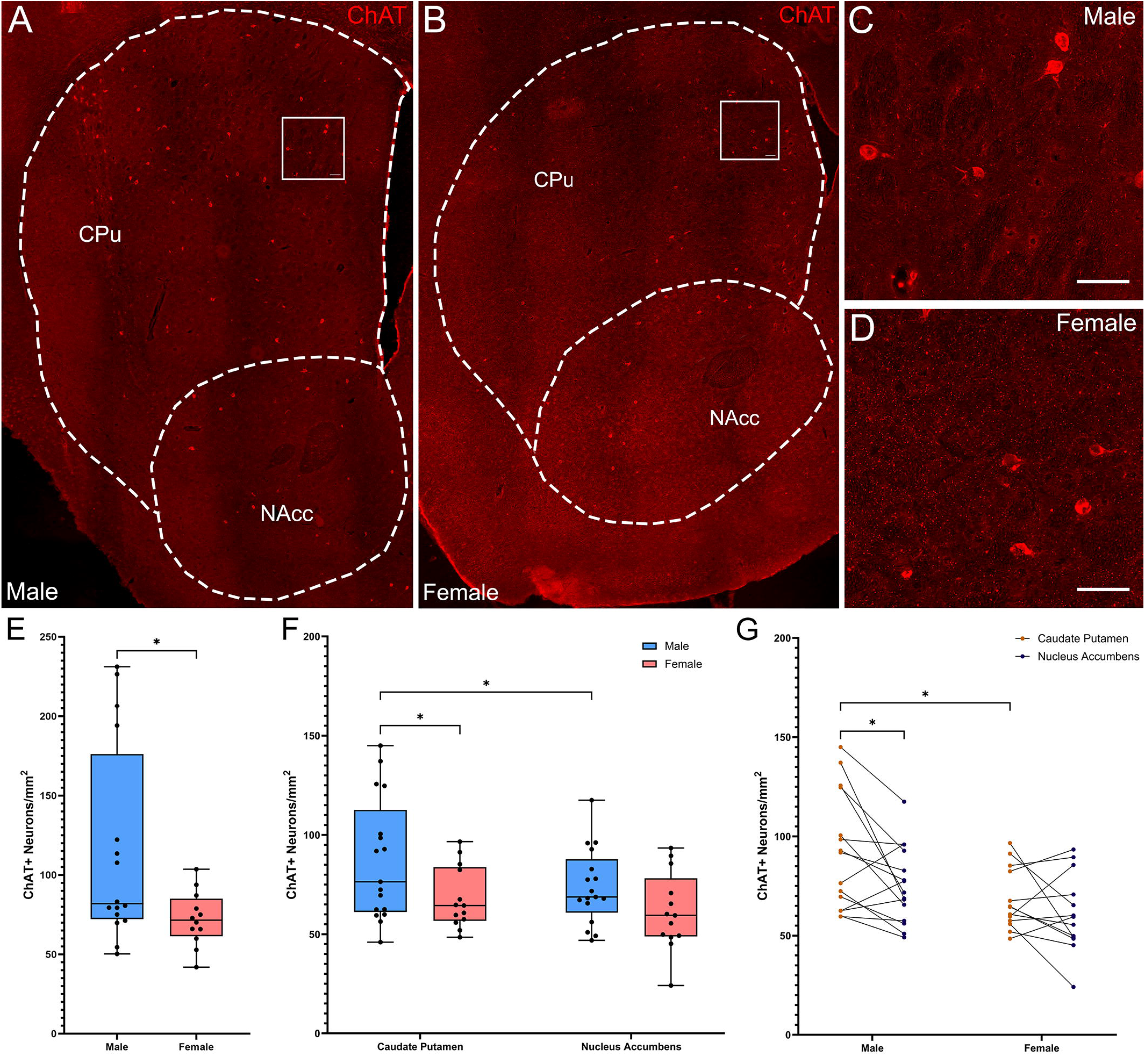
Cholinergic Interneurons are denser and show a dorsally skewed distribution in the striatum of male versus female mice. **(A)** Representative whole striatum reconstruction of high-resolution microscope images of ChAT-expressing interneurons in a male mouse brain, with dashed lines indicating counting regions for CPu and NAcc. **(B)** Representative whole striatum reconstruction of high-resolution microscope images of ChAT-expressing interneurons in a female mouse brain, with dashed lines indicating counting regions for CPu and Nacc. **(C)** Higher resolution image of selected region from A showing CINs in the male CPu. **(D)** Higher resolution image of selected region from B showing CINs in the female CPu. **(E)** Stereological quantification of the density of ChAT+ interneurons per mm^2^ over the whole mouse striatum showing a higher density of ChAT+ neurons in males compared to females (*Two-tailed t-test, t[26] = 2.305, p=0.029). **(F)** Quantification of the density of ChAT+ interneurons divided over the CPu and NAcc sub-regions of the striatum reveals a significant effect of subregion (Repeated Measures ANOVA, f[1,28] = 5.991, p=0.021) and sex (f[1,28] = 4.769, p=0.038); without interaction (f[1,28] = 0.614, p=0.44). Males showed significantly higher ChAT+ interneuron density in the CPu compared to the NAcc (*Fisher’s LSD: p=0.021) while females did not (p=0.278) and there was higher density of ChAT in the male CPu compared to the female CPu (p=0.026). **(G)** Alternate representation of data from F showing density gradients from the CPu to the NAcc indicating a significant gradient from CPu to NAcc in males that is not apparent in females. Scalebars equal 20μm.

To further test for a dorsal-ventral CIN gradient we also analyzed CIN density in dorsal and ventral striatal subregions, defined by an arbitrary line across the middle of the structure (see Methods). This analysis confirmed the differential distribution of CINs in males and females (Supplemental Figure 1A,D-E; Repeated measures ANOVA: subregion x sex interaction: F(1, 24) = 9.012, p = 0.006; effect of subregion: F(1, 24) = 20.42, p<0.001, trend of effect of sex: F(1, 26) = 3.996, p=0.056). CINs exhibited a significant dorsal-ventral gradient in the male striatum (Fisher’s LSD: p<0.001), but not in females (Fisher’s LSD: p=0.311). CINs were denser in male dorsal striatum than in females (Fisher’s LSD: p = 0.006); this pattern was not seen ventrally (p=0.36).

Quantification of overall density of PV+ interneurons in whole striatum (Figure 2A-D) did not reveal significant differences in density between males and females (Figure 2E; t-test, t[36] = 0.595; p=0.56). However, analyzing across subregions (Figure 2A-C) revealed a different distribution in males and females (Figure 2F-G; Repeated Measures ANOVA: main effect of subregion, F(1, 29) = 57.06, p<0.001; sex x subregion interaction, (F (1, 29)=7.745, p=0.009). Both sexes showed significantly higher density of PV+ interneurons in the CPu than in the NAcc (Fisher’s LSD: males p<0.001, females p=0.0013); males showed a significantly higher density of PV+ interneurons in the CPu than females (Fisher’s LSD: p=0.011). Similar effects were seen when cell density was analyzed in arbitrary dorsal and ventral subregions (Supplemental Figure 1B,F,G).

**Figure 2.**
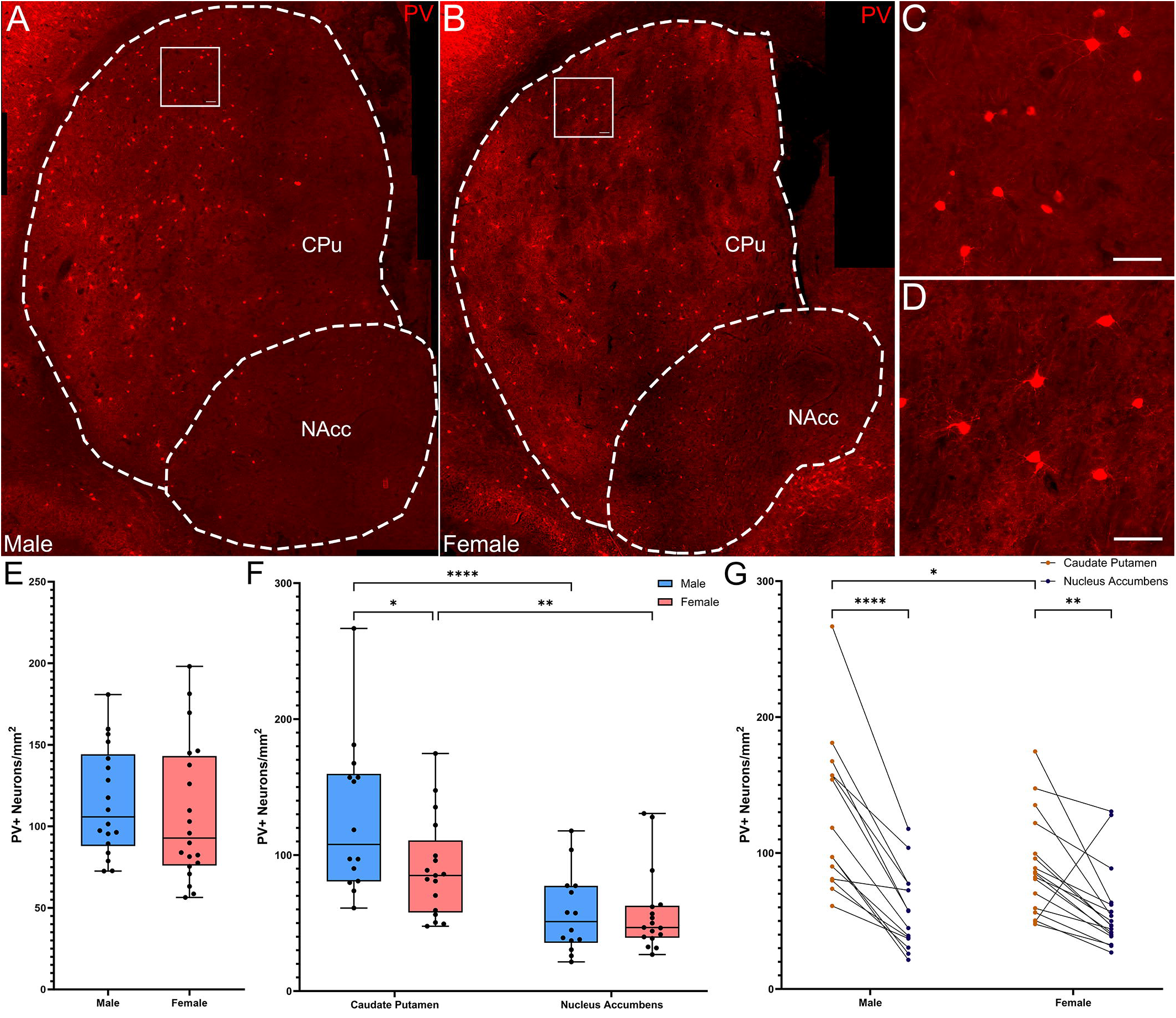
Parvalbumin-expressing interneurons are more concentrated in the dorsal versus the ventral striatum in male, but not female mice. **(A)** Representative whole striatum reconstruction of high-resolution microscope images of PV-expressing interneurons in a male mouse brain, with dashed lines indicating counting regions for CPu and NAcc. **(B)** Representative whole striatum reconstruction of high-resolution microscope images of PV-expressing interneurons in a female mouse brain, with dashed lines indicating counting regions for CPu and Nacc. **(C)** Higher resolution image of selected region from A showing PV+ interneurons in the male CPu. **(D)** Higher resolution image of selected region from B showing PV+ interneurons in the female CPu. **€** Quantification of the density of PV+ interneurons per mm^2^ over the whole mouse striatum using stereological methods shows similar densities of PV+ neurons between sexes (Two-tailed t-test, t[36] = 0.595; p=0.56). **(F)** Quantification of the density of PV+ interneurons divided over the CPu and NAcc sub-regions of the striatum reveals a significant effect of subregion (Repeated Measures ANOVA, f(1, 29) = 57.06, p<0.001) and a significant interaction of sex and subregion (f(1, 29) = 7.745, p=0.009). Males showed significantly higher PV+ interneuron density in the CPu compared to the NAcc (****Fisher’s LSD: p<0.001) while females showed a similar, but less robust gradient (***p=0.001). PV+ interneurons in the CPu were significantly denser in males than in females (*p=0.011), but there were no differences in density in the NAcc (p=0.97). **(G)** Alternate representation of data from F showing density gradients of PV+ interneurons from the CPu to the NAcc. Both males and females showed a dorsal to ventral gradient of PV interneurons, however this was significantly higher in males. Scalebars equal 20μm.

SOM+ interneuron densities across the full striatum did not differ between males and females (Figure 3A-E, t-Test: t[27] = 0.515, p=0.61). Quantification of SOM+ interneurons by subregion (Figure 3F-G) showed a significant effect of subregion (Figure 3F; Repeated Measures ANOVA: F(1, 23) = 27.45, p<0.001), but no effect of sex or subregion x sex interaction. Both males (Fisher’s LSD: p=0.001) and females (Fisher’s LSD: p=0.001) had significantly more SOM interneurons in the NAcc than in the CPu. A similar pattern was seen when cell density was analyzed in arbitrary dorsal and ventral subregions (Supplemental Figure 1C,H,I).

**Figure 3.**
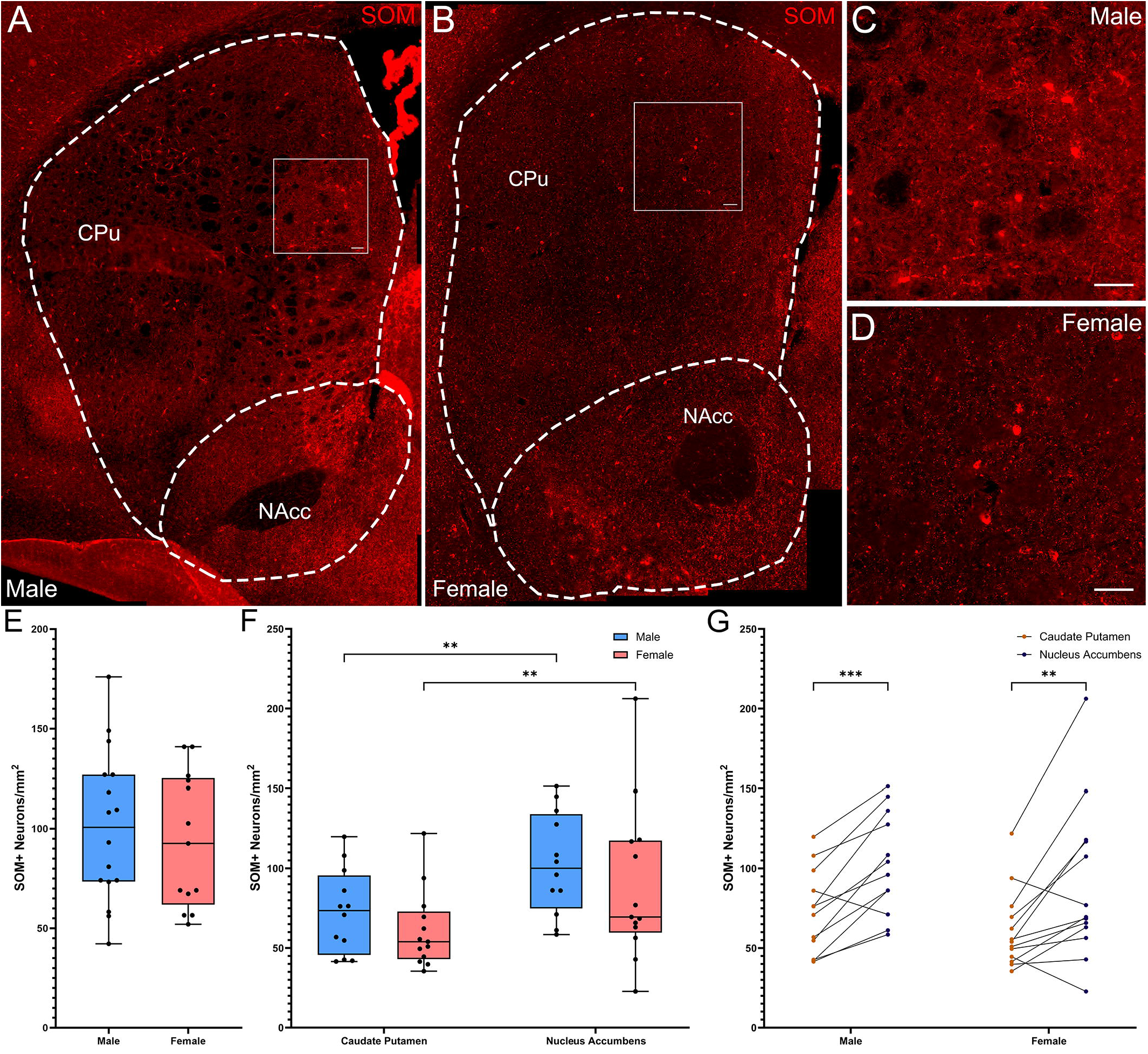
Somatostatin-expressing interneurons are denser in the ventral striatum in both male and female mice. **(A)** Representative whole striatum reconstruction of high-resolution microscope images of SOM-expressing interneurons in a male mouse brain, with dashed lines indicating counting regions for CPu and NAcc. **(B)** Representative whole striatum reconstruction of high-resolution microscope images of SOM-expressing interneurons in a female mouse brain, with dashed lines indicating counting regions for CPu and Nacc. **(C)** Higher resolution image of selected region from A showing SOM+ interneurons in the male CPu. **(D)** Higher resolution image of selected region from B showing SOM+ interneurons in the female CPu. **(E)** Quantification of the density of SOM+ interneurons per mm^2^ over the whole mouse striatum using stereological methods, showing similar densities of SOM+ neurons between sexes (Two-tailed t-test, t[27] = 0.515, p=0.61). **(F)** Quantification of the density of SOM+ interneurons divided over the CPu and NAcc sub-regions of the striatum reveals a significant effect of subregion (Repeated Measures ANOVA, f(1, 23) = 27.45, p<0.001) and no effect of sex (f1, 23) = 0.9600, p=0.337) or interaction (f(1, 23) = 0.01914, p=0.891). Both males and females showed significantly higher SOM+ interneuron density in the NAcc compared to the CPu (**Fisher’s LSD: p=0.001 for both) with no sex differences in density for either the CPu (p=0.404) or the NAcc (p=0.346). **(G)** Alternate representation of the data from F showing a significant density gradient from the NAcc to the CPu in both males and females of equal significance. Scalebars equal 20μm.

## Discussion

Interneurons of the striatum are critical to basal ganglia function (Kreitzer, 2009) and have emerged as a locus of pathology in a number of neuropsychiatric conditions (Poppi et al, 2021; Rapanelli et al, 2017a). Biological sex modulates basal ganglia structure and function (Giedd et al, 1997; Rijpkema et al, 2012), and numerous studies show sex differences in conditions in which basal ganglia dysregulation is implicated, including TS (Martino et al, 2013; Scahill et al, 2014) and autism (Loomes et al, 2017). However, no studies to date have systematically examined sex differences in the distribution, structure, or function of striatal interneurons.

Several populations of striatal interneuron have been found to be reduced in number in post-mortem studies of individuals with persistent adult TS (Kalanithi et al, 2005; Kataoka et al, 2010; Lennington et al, 2016); these studies included both male and female subjects but were not designed to detect sexually dimorphic effects. In mice, we have found experimental depletion of cholinergic interneurons (CINs), parvalbumin-expressing fast-spiking interneurons (FSIs), or both to destabilize the striatum, resulting in repetitive behavioral pathology, social deficits, and other behavioral abnormalities (Xu et al, 2015; Xu et al, 2016; Rapanelli et al, 2017; Cadeddu et al, 2023). Interestingly, these effects are sexually dimorphic, appearing in male but not female mice, despite equivalent levels of interneuron depletion (Rapanelli et al, 2017; Cadeddu et al, 2023). The underlying differences in striatal anatomy or function that lead to this differential susceptibility remain unclear.

Here, we document intriguing differences between male and female adult mice in striatal interneuron density and distribution. Differential density and distribution of interneurons suggests differences in striatal function, and perhaps in resilience and vulnerability to pathophysiological processes.

Cholinergic interneurons (CINs) play a central role in the striatal microcircuitry. CINs regulate the release of both dopamine (Cachope et al, 2012; Threlfell et al, 2012) and GABA (Nelson et al, 2014) from dopaminergic afferents from the substantia nigra. They also modulate multiple populations of GABAergic interneurons, as well as MSNs (Poppi et al, 2021). We find a significant difference between sexes in both density and distribution of these interneurons: The density of CINs is significantly higher in the CPu in males than in females. The existence of a dorsal to ventral gradient of CINs in males is consistent with previous findings in males (Matamales et al, 2016; Phelps et al, 1989); CIN density in females has not previously been rigorously quantified. Our findings match observations in previous studies, in which a higher CIN density was observed in males than in females (Rapanelli et al, 2017; Cadeddu et al, 2023); however, these previous observations were incidental, without the careful quantification applied here. While female mice have relatively even distribution of ChAT+ interneurons between the dorsal and ventral striatum, males a greater density in the CPu compared to the NAcc.

The presence of more CINs in the dorsal striatum of males than of females is perhaps surprising, as our previous work shows that males are more sensitive to disruption of these cells (Rapanelli et al, 2017; Cadeddu et al, 2023). It might have been expected that a higher basal density of CINs would render males more resilient to CIN depletion. The current data suggest a different model, that the dorsal regulatory circuitry in males may be more reliant on CINs than that of females and thus is less able to accommodate pathological CIN disruption.

PV-expressing FSIs are highly active interneurons innervating a wide variety of targets in the striatum through both inhibition and disinhibition circuits (Duhne et al, 2021), and are particularly important in regulating MSNs through feedforward inhibition (Adler et al, 2013; Gittis et al, 2011; Gittis et al, 2010; Koós & Tepper, 1999; Lee et al, 2017). There is previous evidence of a dorsoventral gradient of PV distribution (Zahm et al, 2003) in addition to a rostrocaudal gradient (Wu & Parent, 2000); sex effects have not not previously been examined. We find effects of both sex and striatal subregion on FSI density. Male mice again exhibit a significantly higher density in the dorsal CPu than in the ventral NAcc; females also show a difference in density in the same direction, but it is less robust. Because FSIs are an important source of inhibition in the striatum, this may suggest that the male and female striatum differ in their excitatory-inhibitory balance. We have previously found depletion of FSIs in the dorsal striatum to produce both repetitive behavioral pathology and elevated anxiety-like behavior (Xu et al, 2016), though it is not yet clear if these effects are specific to males. When both FSIs and CINs are depleted, behavioral effects are seen only in males, suggesting that there may be a sex-specific effect (Rapanelli et al, 2017b).

FSIs and CINs separately regulate MSN activity (Adler et al, 2013; Gritton et al, 2019), and each other (Chang & Kita, 1992; DeBoer & Westerink, 1994; Faust et al, 2016). Sex differences in density and distribution follows the same general pattern for both interneuron types, with higher density in the dorsal CPu region of the striatum and a greater dorsal-ventral gradient towards the NAcc in males than in females in both cases. This may suggest a single developmental process underlying both effects.

We found no sex differences in the density or distribution of SOM-expressing GABAergic interneurons. However, both males and females had more SOM-expressing interneurons in the NAcc compared to the CPu. This pattern, quite distinct from that seen with CINs and FSIs, indicates that the sex differences are not uniform across all components of the striatal microcircuitry but rather differentially affect distinct cells and processes. This reinforces the idea that differential density, distribution, and function of interneurons in males and females leads to distinct regional differences in information processing in the basal ganglia circuitry.

This study has several limitations that should be addressed in future work. First, while we used stereological tools to ensure that we did not overcount cells in the examined striatal slices, we did not systematically reconstruct the full anterior-posterior extent of the striatum, and thus we cannot comment on anterior-posterior gradients or calculate the total number of interneurons in the male or female striatum, or its subregions. Second, we rely on the expression of specific markers to identify interneurons; our counts are thus susceptible to error if these markers are differentially expressed in males and females. That said, staining was strong and was qualitatively similar in males and females (see representative micrographs in all figures). Finally, we have examined interneuron number but not more granular aspects of their structure, function, or integration into local microcircuitry. These are important directions for future studies.

In sum, we find sex differences in the density and distribution of striatal interneurons, with a higher density of both CINs and FSIs in the dorsal striatum in males. The mechanisms underlying these differential patterns of interneuron density and distribution remain to be elucidated but are likely to involve sexually dimorphic modulation of developmental processes. Of note, estrogen receptors are expressed on both cholinergic (Almey et al, 2012; Kövesdi et al, 2022) and GABAergic interneurons (Almey et al, 2016), providing a potential mechanism for sex hormones to regulate interneuron function. These findings represent a starting point for future work analyzing the impact of these differences on sex dependent outcomes in both normal basal ganglia function and in the pathophysiology of a range of neuropsychiatric conditions, such as TS and ASD.

## Author Contributions

MVZ: Conceptualization, Formal Analysis, Funding Acquisition, Investigation, Project Administration, Visualization, Writing – Original Draft

DF: Investigation

CP: Conceptualization, Funding Acquisition, Project Administration, Resources, Supervision, Writing – Review and Editing

## Acknowledgments

The authors thank Betsy D’Amico and the staff of the Yale Animal Resource Core and Charles River Laboratories for animal husbandry.

