## Supplemental Figure 1 for "Sex differences in the distribution and density of regulatory interneurons in the striatum"

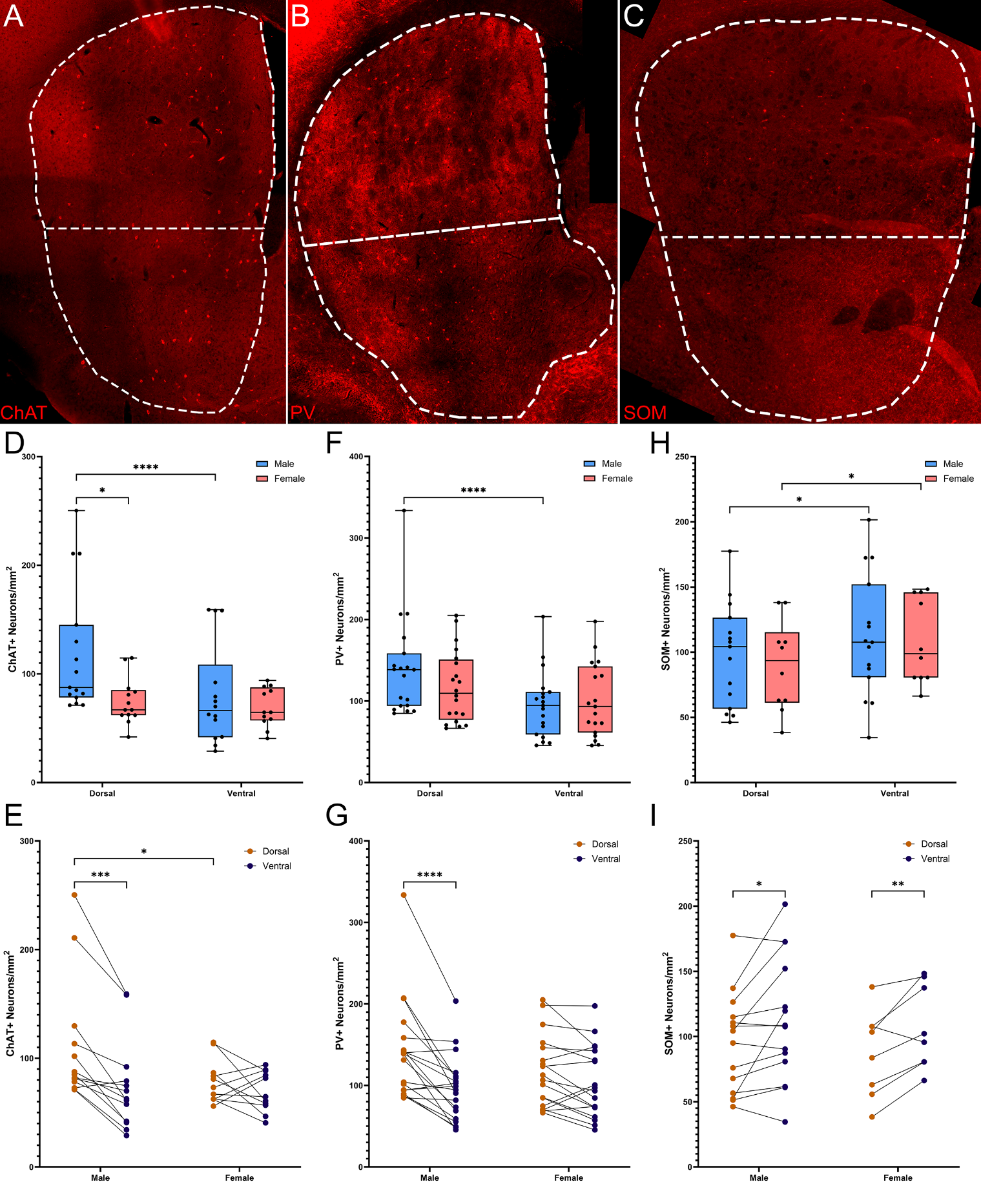

**Supplementary Figure 1.** **Dorsal to ventral quantification of striatal interneurons show sex differences in distribution. (A)** Representative whole striatum reconstruction of high-resolution microscope images of ChAT-expressing interneurons from mouse striatum. Dashed lines indicate counting regions for dorsal and ventral striatum. **(B)** Representative whole striatum reconstruction of high-resolution microscope images of PV-expressing interneurons from mouse striatum. Dashed lines indicate counting regions for dorsal and ventral striatum. **(C)** Representative whole striatum reconstruction of high-resolution microscope images of SOM-expressing interneurons from mouse striatum. Dashed lines indicate counting regions for dorsal and ventral striatum. **(D)** Quantification of the density of ChAT+ interneurons divided over the dorsal and ventral sub-regions of the striatum reveals a significant effect of subregion (f(1, 24) = 20.42, p<0.001), a trend towards significant effect of sex (f(1, 26) = 3.996, p=0.056), and a significant interaction of subregion and sex (Repeated Measures ANOVA, f(1, 24) = 9.012, p=0.006). ChAT+ interneurons in the male dorsal striatum were significantly denser than both female dorsal striatum (*Fisher’s LSD, p=0.005) and male ventral striatum (**p<0.001). **(E)** Alternative representation of data from D showing a significant density gradient from the dorsal to ventral striatum in males, but not females. **(F)** Quantification of the density of PV+ interneurons divided over the dorsal and ventral sub-regions of the striatum reveals a significant effect of subregion (F(1, 35) = 26.91, p<0.001) and a significant interaction of subregion x sex (Repeated Measures ANOVA, f1, 35) = 7.726, p=0.009). Males show a different distribution of PV+ interneurons than females, in that they are significantly denser in the dorsal striatum (****Fisher’s LSD, p<0.001), while females had a more equal distribution between the dorsal and ventral regions (p=0.10). **(G)** Alternative representation of data from F showing a significant dorsal to ventral density gradient of PV+ interneurons in males but not females. **(H))** Quantification of the density of SOM+ interneurons divided over the dorsal and ventral sub-regions of the striatum reveals an effect of subregion (f[1,23] = 9.928, p=0.004) but no effect of (f(1, 23) = 0.1389, p=0.71) or interaction with sex (f(1, 23) = 0.1956, p=0.663). **(I)** Alternate representation of data from H showing ventral to dorsal gradients of SOM+ interneurons in both males and females with no sex difference.
